# Comparative Transcriptomics Reveals Genes Commonly Induced by Distinct Stressors in *Chlamydia*

**DOI:** 10.64898/2025.12.30.696969

**Authors:** Ronald Haines, Danny Wan, Guangming Zhong, Huizhou Fan

## Abstract

*Chlamydia trachomatis* is a leading cause of urogenital infections that can result in serious long-term complications. This obligate intracellular bacterium undergoes a biphasic developmental cycle alternating between the infectious elementary body and the replicative reticulate body and can enter a persistent state in response to adverse environmental conditions. Although transcriptomic reprogramming is central to chlamydial stress adaptation and persistence, how responses differ across biologically distinct stressors remains incompletely defined. Here, we performed a comparative reanalysis of five published, high-quality *C. trachomatis* RNA-Seq datasets generated under prolonged interferon-γ treatment, tryptophan starvation, iron starvation, penicillin exposure, or acute heat shock. Global transcriptomic analyses reveal extensive stress-specific reprogramming and a clear separation between the transcriptome induced by heat shock and those induced by chronic stresses. Transcriptomic overlap observed among chronic stress conditions is substantially reduced when the heat shock transcriptome is included, indicating that shared transcriptional features are stressor-dependent. Unexpectedly, tryptophan starvation and iron starvation exhibit particularly close transcriptomic similarity, consistent with regulatory cross-talk mediated by the iron-dependent transcriptional repressor YtgR. In contrast, interferon-γ induces a distinct but partially overlapping transcriptome, likely reflecting activation of additional host-mediated antimicrobial mechanisms beyond tryptophan deprivation. Together, these findings demonstrate that adaptation to different biological stressors in *C. trachomatis* is driven by distinct transcriptomic reprogramming, while consistently involving a subset of functions that may represent points of vulnerability for disrupting chlamydial persistence.

## INTRODUCTION

Bacteria have evolved strategies to respond to environmental challenges, including nutrient deprivation, sudden temperature fluctuations, and exposure to antibiotics. These adaptive responses are critical for bacteria to survive and thrive in hostile environments (1–3). Pathogenic bacteria, in particular, face additional pressures from host immune defenses and often deploy mechanisms to evade these responses (3). Understanding how bacteria reprogram their cellular functions in response to stress is crucial for deciphering their survival strategies and may aid in the development of novel therapies.

*Chlamydia trachomatis* is an obligate intracellular bacterial pathogen known for causing pelvic inflammatory disease, infertility, ectopic pregnancy, and abortion in women (4–6). Increasing evidence also implicates *C. trachomatis* infection as a contributor to male infertility (7, 8). In addition, ocular serovars of *C. trachomatis* cause trachoma, a major cause of preventable blindness in resource-limited regions worldwide.

*Chlamydia* has a developmental cycle that alternates between the infectious elementary body (EB) and the proliferative reticulate body (RB) (4, 6, 9, 10). EBs enter host cells and differentiate into RBs inside cytoplasmic vacuoles known as inclusions. Following several rounds of replication, RBs differentiate back into EBs, which exit host cells through either host cell rupture or inclusion extrusion (4, 6, 9, 10).

Adverse environmental conditions can disrupt this developmental cycle and induce a persistent state characterized by the formation of enlarged, non-dividing RBs, often referred to as aberrant bodies (11–20). During persistence, RB replication and EB production are suppressed, allowing chlamydiae to survive prolonged stress. Importantly, persistence is reversible; upon stress removal, aberrant bodies revert to RBs, which resume replication and re-enter the productive developmental cycle. Persistence is a major cause of clinical treatment failure and chronic chlamydial infections, contributing to infertility and other complications (11–20).

Chlamydial persistence has been modeled in cell culture using a variety of stressors relevant to infection and therapy, including interferon-γ (IFNγ), β-lactam antibiotics, nutrient deprivation, iron starvation, and heat shock (11, 13–18, 21–24). IFNγ, produced by activated immune cells, inhibits RB replication and EB formation (11, 14–17, 22). β-Lactam antibiotics such as amoxicillin, which may be administered during pregnancy or co-infection, are potent inducers of persistence despite not being a first-line antichlamydial agent (14–17, 23). IFNγ, produced by T-lymphocytes and other immune cells, inhibits RB division and EB production (11, 14–17, 22, 25). Anemia and other conditions that limit nutrient or iron availability can restrict chlamydial growth (26–28). Febrile responses are more characteristic of infections caused by *C. trachomatis* lymphogranuloma venereum serovars or the zoonotic pathogen *Chlamydia psittaci* (29–33) and may expose bacteria to repeated episodes of elevated temperatures that disrupt protein homeostasis (21, 34, 35).

Stress responses in *C. trachomatis* are governed by transcriptomic reprogramming (11, 13, 16, 21, 24). Previous studies have shown overlaps in transcriptomic changes induced by specific stressors, such as iron starvation and tryptophan depletion (24) or β-lactam antibiotics and IFNγ (16). However, how transcriptional responses compare across a broader range of chronic and acute stress conditions, and the extent to which similarities or differences reflect shared or distinct challenges, remain incompletely defined (21).

To address this gap, we performed a comparative reanalysis of published *C. trachomatis* RNA-Seq datasets generated under five stress conditions: IFNγ treatment (16), iron starvation (24), tryptophan starvation (24), penicillin exposure (16), and heat shock (21). By systematically comparing these transcriptomes using a unified analytical approach, this study demonstrates that adaptation of *C. trachomatis* to diverse biological stressors is driven by distinct transcriptional programs. It further establishes that a set of regulated genes is consistently involved across stress conditions and may represent shared vulnerabilities that may be exploited for therapeutic intervention.

## RESULTS

### Exceptionally high sequencing depths of stress transcriptomic studies

We analyzed published RNA-Seq datasets from *C. trachomatis* cultures exposed to five distinct stress conditions: the antichlamydial cytokine IFNγ, the β-lactam antibiotic penicillin, iron starvation induced by the chelator 2,2-bipyridyl, tryptophan starvation using a tryptophan-free medium, and heat shock (16, 21, 24). Key experimental parameters are summarized in Table 1. The IFNγ and penicillin stress studies utilized *C. trachomatis* serovar D (16), whereas the remaining studies employed serovar L2 (21, 24). Human cervical carcinoma HeLa cells served as the host cell line in all experiments except for the heat shock study, which used mouse fibroblast L929 cells (16, 21, 24).

**Table 1.** Experimental conditions, dataset sources, and depths for the stress transcriptomic studies analyzed. For IFNγ and penicillin treatments, duplicate biological samples were sequenced separately and deposited under distinct accession numbers. For iron and tryptophan starvation studies, triplicate biological samples were sequenced and deposited under a single accession number. Abbreviations: Ct, *C. trachomatis*; hpi, hours postinoculation; BPD, 2,2-bipyridyl.

| Study | Ct organism | Host cell | Treatment & duration | Raw RNA-Seq data accession # | Mapped Ct reads | Depth (x coverage) |
| --- | --- | --- | --- | --- | --- | --- |
| Interferon-γ (16) | Serovar D (UW-3/Cx) | HeLa | Control medium | SRR5834397,<br>SRR5834399, | 16800722<br>9755404 | 1600<br>929 |
| | | | 50 U/mL IFN-γ,<br>-24 to 24 hpi | SRR13189638,<br>SRR13189738 | 2557223<br>4331636 | 244<br>413 |
| Penicillin (16) | Serovar D UW-3/Cx | HeLa | Control medium | SRR5834397,<br>SRR5834399 | 16800722<br>9755404 | 1600<br>929 |
|  |  |  | 1 U/mL penicillin,<br>0 to 24 hpi | SRR5834398,<br>SRR5834400 | 31724401<br>18660673 | 3021<br>1777 |
| Iron starvation (24) | Serovar L2 (434/BU) | HeLa | Control medium | GSE179003 | 118754751 | 8512 |
| | | | 100 μM BPD,<br>0 to 24 hpi | GSE179003 | 42374135 | 3037 |
| Tryptophan starvation (24) | Serovar L2 (434/BU) | HeLa | Control medium | GSE179003 | 118754751 | 8512 |
|  |  |  | Trp-free medium,<br>0 to 24 hpi | GSE179003 | 16509535 | 1183 |
| Heat shock (21) | Serovar L2 (434/BU) | L929 | 37 °C | GSE173366 | 5,435,469 | 260 |
|  |  |  | 45 °C,<br>15.5 to 16 hpi | GSE173366 | 5,681,297 | 271 |

Tryptophan starvation, iron starvation, and penicillin treatment were applied after inoculation, whereas IFNγ treatment of host cells was initiated 24 h prior to inoculation. Tryptophan- or iron-starved and IFNγ- or penicillin-treated cultures were harvested at 24 h postinoculation for RNA extraction. Heat shock treatment at 45 °C was conducted between 15.5 and 16 h postinoculation (21). Based on the length of stress exposure, we characterize the first four transcriptomes as chronic stress transcriptomes and heat shock as an acute stress transcriptome.

To ensure comparability across studies, we reprocessed all raw sequencing data using a consistent bioinformatic workflow and identical software tools, rather than relying on previously published secondary analyses. All five RNA-Seq datasets achieved extremely high sequencing depths (Table 1), with genome coverage ranging from nearly 250-fold to over 8,000-fold. These high coverages support rigorous comparisons of *Chlamydia* transcriptomic responses across diverse stress conditions.

### Extensive transcriptomic reprogramming in stress responses

The *C. trachomatis* genome encodes more than 900 protein-coding genes. DESeq analysis revealed that exposure to IFNγ, tryptophan starvation, iron starvation, penicillin, or heat shock markedly altered the expression of nearly one-third of these genes, ranging from 292 to 363 differentially expressed genes (DEGs) at a ≥1.5-fold change and *P* < 0.05 threshold (Table 2). These findings reinforce previous observations that *C. trachomatis* mounts extensive transcriptomic reprogramming when confronted with diverse forms of stress (11, 13, 16, 21, 24).

**Table 2.** Numbers transcriptome. of upregulated and downregulated genes within each stress.

| **Stressor** | **Upregulated genes** | **Downregulated genes** | **All DEGs** |
| --- | --- | --- | --- |
| Interferon-γ | 163 | 200 | 363 |
| Iron Starvation | 136 | 181 | 317 |
| Tryptophan Starvation | 154 | 189 | 343 |
| Penicillin | 92 | 200 | 292 |
| Heat Shock | 171 | 156 | 325 |

### Distinction of IFNγ, tryptophan starvation, iron starvation, or penicillin stress transcriptomes from the heat shock transcriptome

To further compare the five stress transcriptomes, we generated a heatmap for the 667 genes that were differentially expressed in at least one condition (Fig. 1A). Hierarchical clustering revealed that tryptophan starvation and iron starvation formed the closest pair, reflecting their broadly similar effects on the bacterium due to nutrient starvation. Penicillin-treated cultures clustered next to this pair. In contrast, the IFNγ transcriptome was positioned farther away from the tryptophan-starvation transcriptome, an unexpected separation given the well-established link between IFNγ exposure and host-driven tryptophan starvation. The heat shock transcriptome remained the most distinct, forming a well-separated branch relative to the four chronic stress conditions.

**Fig. 1.**
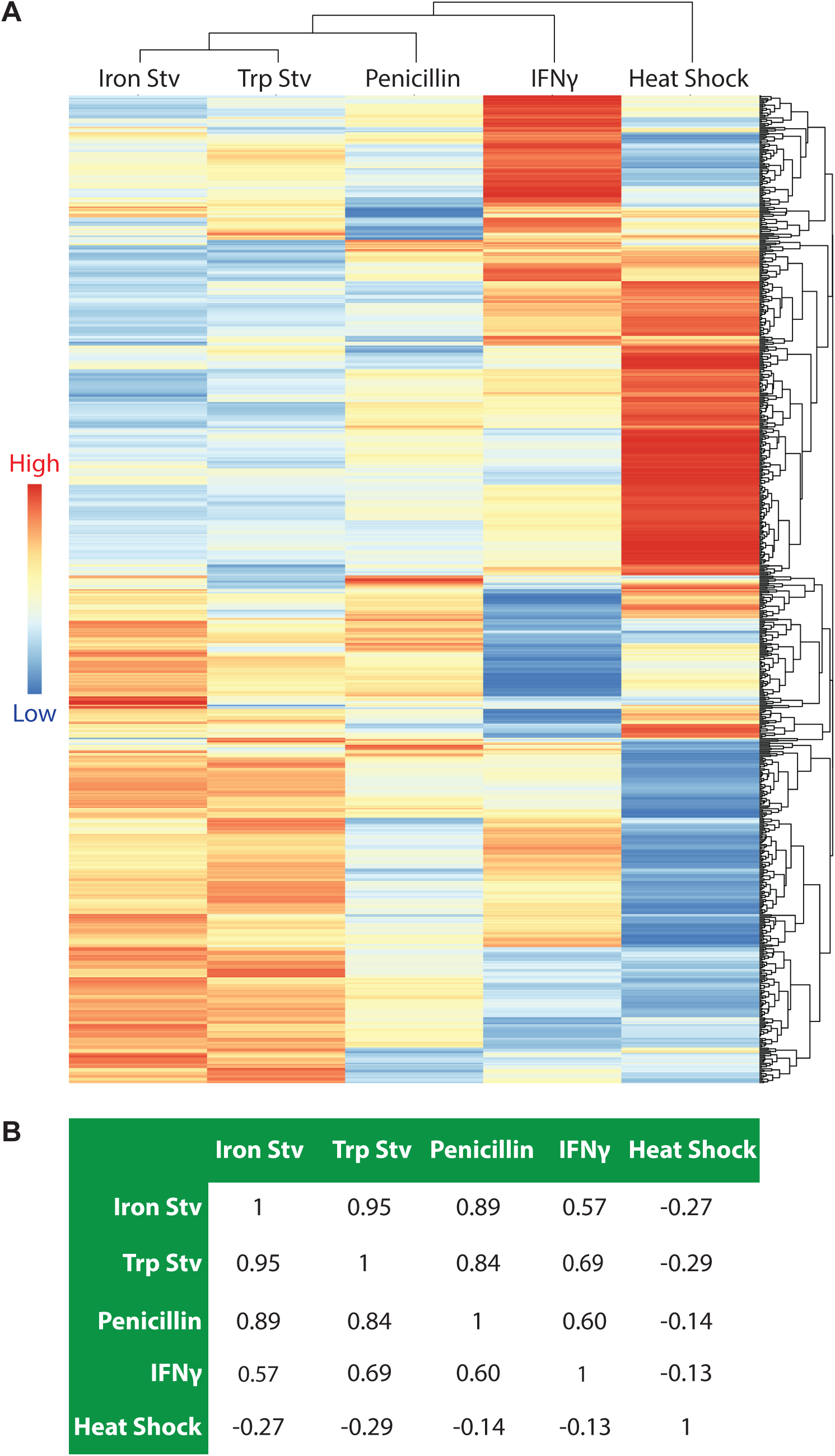
Acute heat shock and chronic stressors elicit distinct transcriptomic responses in *C. trachomatis*. (A) Hierarchical clustering heatmap of 667 genes differentially expressed in at least one stress transcriptome (adjusted *P* < 0.05; |log_2_ fold change| ≥ 0.585). Each row represents a gene, and each column represents a stressor. The dendrograms depict the relative similarity among genes (rows) and among stress transcriptomes (columns) based on their overall expression patterns across conditions. Heatmap colors represent log₂ fold change values relative to the corresponding control condition. The heatmap visualization was generated using *pheatmap*. (B) Pairwise Pearson correlation analysis of stress transcriptomes based on log₂ fold-change values of the same gene set, providing an independent quantitative measure of transcriptome similarity. Abbreviations: Stv, starvation; Trp, tryptophan.

To independently assess the relatedness among the five transcriptomes, we performed pairwise Pearson correlation analysis using the log₂ fold-change values for the aforementioned 667 DEGs (Fig. 1B). Tryptophan starvation and iron starvation again showed the strongest correlation (r = 0.95). Penicillin treatment correlated strongly with iron starvation (r = 0.89) and tryptophan starvation (r = 0.84). The IFNγ transcriptome displayed more moderate correlations with these three conditions (r = 0.57-0.69), consistent with its more distant placement in the clustering analysis (Fig. 1A). Heat shock exhibited negative correlations with all other conditions (r = -0.14 to -0.29), reinforcing its distinct transcriptomic profile. Together, hierarchical clustering and correlation analysis reveal a clear reprogramming pattern among the five stress transcriptomes, with nutrient starvation- and penicillin-induced transcriptomes forming a closely related group, IFNγ forming a more separate branch, and heat shock representing the most divergent condition.

### Opposing effects of acute and chronic stress on the expression of protein synthesis genes and type III secretion system genes

To examine how *C. trachomatis* prioritizes biological processes under different stress conditions, we assigned the DEGs to functional categories and displayed the distributions of up-and down-regulated genes as pie graphs (Fig. 2). In the IFNγ, tryptophan starvation, iron starvation, and penicillin transcriptomes, the translation and ribosomal structure and biogenesis category accounted for the largest fraction of upregulated genes, representing 23-37% of all upregulated genes under these chronic stresses. In contrast, this category contributed only 3-7% of the downregulated genes in the chronic stress transcriptomes but represented 25% of all downregulated genes under heat shock. Thus, translation and ribosomal structure and biogenesis category genes were preferentially upregulated during chronic stress but downregulated during acute heat shock.

**Fig. 2.**
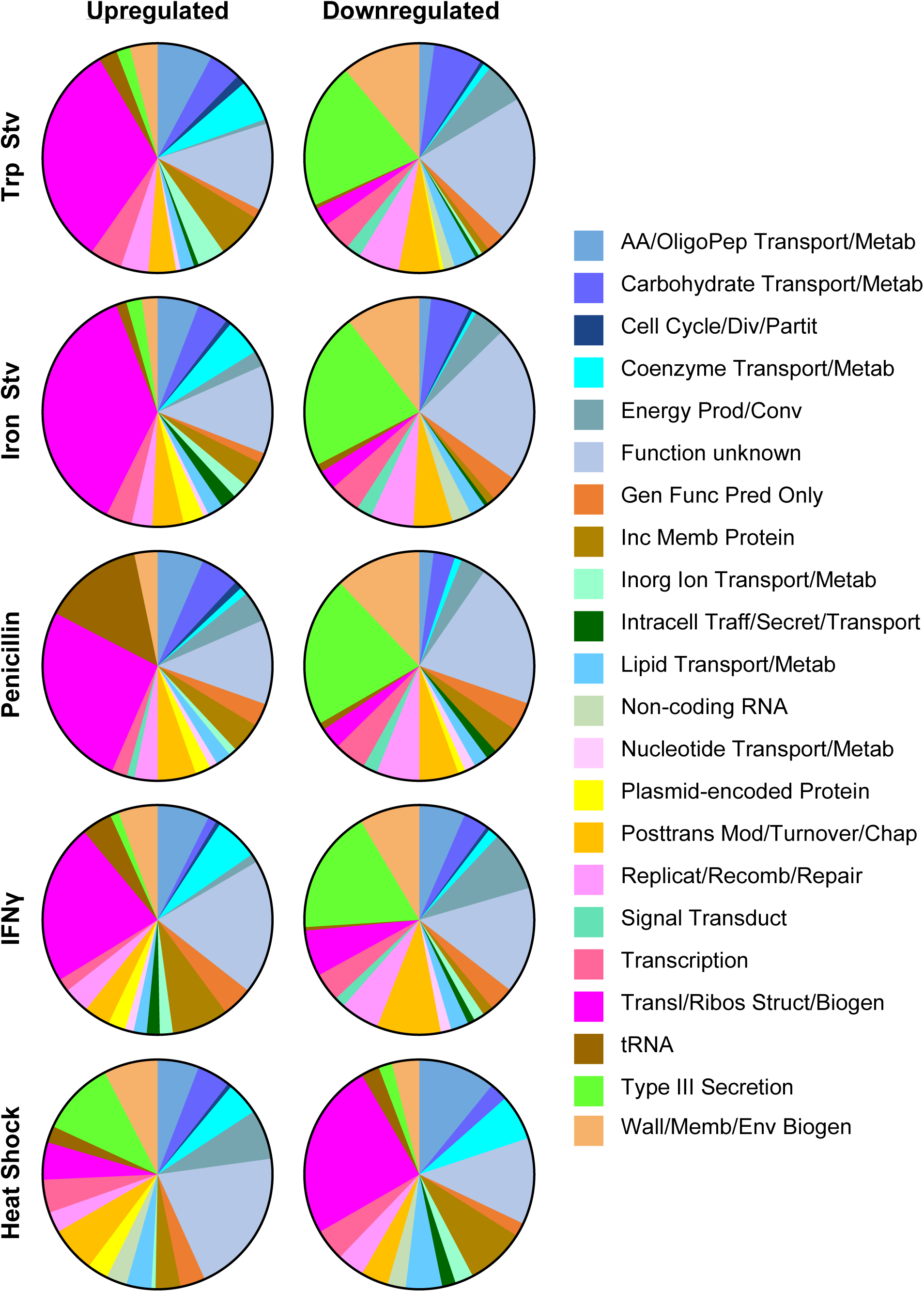
Acute and chronic stressors exert opposing effects on protein synthesis and T3SS gene expression. Pie charts show the distribution of upregulated (left) and downregulated (right) genes among gene ontology categories within each stress transcriptome. Only genes meeting differential expression criteria (adjusted *P* < 0.05; |log₂ fold change ≥ 0.585) are included. Excluding genes of unknown function, genes involved in translation and ribosomal structure and biogenesis dominate the upregulated gene sets in chronic stress transcriptomes but the downregulated gene sets in the heat shock transcriptome, whereas T3SS genes dominate the downregulated gene sets in chronic stress transcriptomes but the upregulated gene set in the heat shock transcriptome. Abbreviations: AA, amino acid; OligoPep, oligopeptide; Metab, metabolism; Div, division; Partit, partitioning; Prod/Conv, production and conversion; Gen Func Pred, general function prediction; Inc Memb, inclusion membrane; Inorg, inorganic; Traff, trafficking; Secret, secretion; Posttrans Mod, posttranslational modification; Chap, chaperone; Replicat, replication; Recomb, recombination; Transduct, transduction; Wall/Memb/Env Biogen, wall, membrane, or envelope biogenesis.

A reciprocal pattern was observed for the type III secretion system (T3SS) category. Under the four chronic stress conditions, T3SS genes constituted only 1-3% of upregulated genes but comprised 18-22% of downregulated genes. Specifically, *copD*, *scc2*, *ctl0003/ct635*, *ctl0063/ct694*, *ctl0080/ct711*, *ctl0081/ct712*, *ctl0219/ct847*, *ctl0220/ct843*, *ctl0255/ct875*, *ctl0338/ct082*, *ctl0338A/ct083*, *ctl0397/ct142*, *ctl0398/ct143*, *ctl0883/ct619*, and *ctl0884/ct620*, which encode T3SS structural components, chaperones, or effectors, were downregulated in all four chronic stress transcriptomes (Dataset S1). Previous studies have shown that these genes are upregulated during RB-to-EB differentiation (36–38). The predominant downregulation of T3SS genes during chronic stress is consistent with the persistent state, in which RB-to-EB differentiation and late developmental gene expression are suppressed. Several type III-secreted effectors participate in late developmental events linked to chlamydial exit, including Inc proteins that regulate inclusion extrusion (e.g., CT228 and MrcA) and the effector CteG, which promotes host cell lytic exit (39–43). Their downregulation under chronic stress likely helps retain persistent chlamydiae intracellularly rather than promoting their release. In contrast, heat shock at 45 °C upregulated 11% of its total upregulated genes in the T3SS category while downregulating only 2% of its downregulated genes (21), suggesting that T3SS effectors normally induced during late development may contribute to intracellular chlamydial survival under this extreme stress condition.

### Stress-specific distinctions among transcriptomes

To complement the DEG count-based analysis shown in the pie graphs (Fig. 2), we calculated the fraction of genes that were upregulated or downregulated within each functional category and displayed these values as divergent bar graphs (Fig. 3; Fig. S1). This category-normalized approach confirms the opposing behaviors of translation, ribosomal structure and biogenesis, and T3SS genes described above and further distinguishes the acute heat shock transcriptome from those induced by chronic stress conditions (Fig. S1). Importantly, this analysis also reveals regulatory patterns that are not apparent from comparisons based solely on absolute DEG counts.

**Fig. 3.**
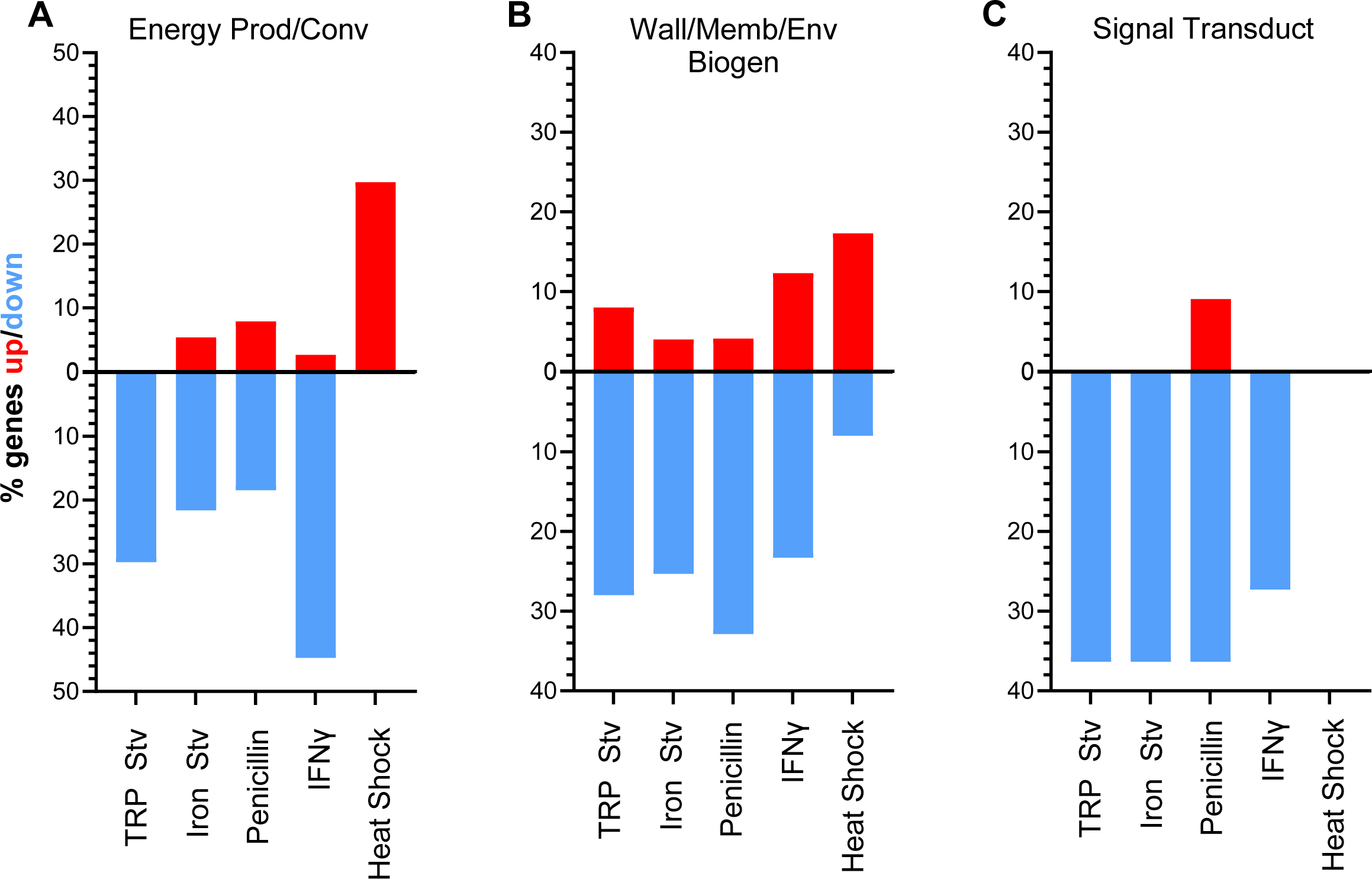
Category-normalized analysis highlights stress-specific regulation of functional gene groups. Divergent bar plots show the fraction of genes that are upregulated (red) or downregulated (blue) within selected gene ontology categories across five stress transcriptomes. Values represent the percentage of genes within each category that meet the differential expression criteria under each condition, thereby normalizing for differences in category size. Categories shown are energy production and conversion (A), wall, membrane, or envelope biogenesis (B), and signal transduction mechanisms (C).

Three functional categories displayed particularly strong differences between the chronic stress transcriptomes and the acute heat shock transcriptome, evident in both the upward and downward components of the divergent bars. Genes involved in energy production and conversion showed only small upward fractions and large downward fractions across all four chronic stress transcriptomes, whereas the heat shock transcriptome exhibited a prominent upward fraction with virtually no corresponding downregulation (Fig. 3A). This opposing pattern is consistent with reduced growth and metabolic activity under chronic stress, which likely lowers cellular energy demand, whereas survival under acute heat shock requires enhanced energy production to support stress tolerance and recovery.

Genes involved in cell wall, membrane, or envelope biogenesis exhibited consistently large downward fractions and minimal upward fractions across all four chronic stress transcriptomes, whereas the heat shock transcriptome showed a noticeable upward fraction with little corresponding downregulation (Fig. 3B). Many of the genes downregulated under chronic stress are preferentially expressed in EBs. Notably, *omcA* and *omcB*, which encode major components of the EB outer membrane complex (44), were downregulated in all four chronic stress transcriptomes but upregulated during heat shock (Dataset S1). This pattern is consistent with suppression of EB-associated envelope biogenesis during chronic stress and reflects inhibition of RB-to-EB differentiation under persistence-inducing conditions. In contrast, the upregulation of *omcA* and *omcB* during acute heat shock is unexpected and may reflect a distinct stress response, possibly utilizing EB envelope components to help maintain chlamydial cellular integrity under extreme conditions.

Signal transduction mechanism genes also exhibited a clear pattern at the functional category level in the fractional analysis (Fig. 3C; Fig. S1). Across all four chronic stress conditions, this category showed a strong net downward bias, whereas heat shock induced little overall change. However, this coordinated behavior at the category level did not reflect consistent regulation of individual signaling components. For example, expression of the anti-sigma factor gene *rsbW* remained largely unchanged across stress conditions, whereas the anti-anti-sigma factor gene *rsbV2* displayed stress-dependent regulation (Dataset S1). These findings suggest that signal transduction pathways are reprogrammed through selective and condition-specific regulatory adjustments across stress conditions.

Beyond this category-level trend, the fractional analysis revealed pronounced stress-specific regulation in several functional categories (Fig. S1). For example, inorganic ion transport and metabolism exhibited prominent upregulation during tryptophan starvation and, to a lesser extent, IFNγ treatment, but showed little response under iron starvation or exposure to penicillin. In contrast, tRNA genes exhibited highly variable upregulation across chronic stresses, with strong induction during penicillin treatment but minimal changes under iron starvation. These patterns underscore that, alongside common features, each stress condition elicits a distinct transcriptional response rather than a common chronic stress program.

Notably, the plasmid-encoded virulence gene *pgp3*, which encodes the secreted effector Pgp3 (45–48), and *pgp4*, which encodes a transcriptional regulator of pgp3 and numerous chromosomal genes (49, 50), exhibited stress-dependent regulation (Fig. S1). *pgp3* was downregulated during tryptophan starvation and penicillin exposure, whereas *pgp4* was repressed during penicillin treatment but induced during heat shock (Dataset S1). The other plasmid genes, which primarily support plasmid maintenance, also showed stress-dependent regulation, but without a consistent pattern across conditions (Dataset S1).

### Expression patterns of transcriptional regulator genes across stress transcriptomes

To examine how transcriptional regulation contributes to the organization of the five stress transcriptomes, we analyzed the expression patterns of transcriptional regulator genes that were differentially expressed in at least one condition and visualized their profiles by hierarchical clustering (Fig. 4). Clustering based on regulator expression closely mirrored clustering of the global transcriptomes, reinforcing the clear separation between the acute heat shock response and the four chronic stress responses.

**Fig. 4.**
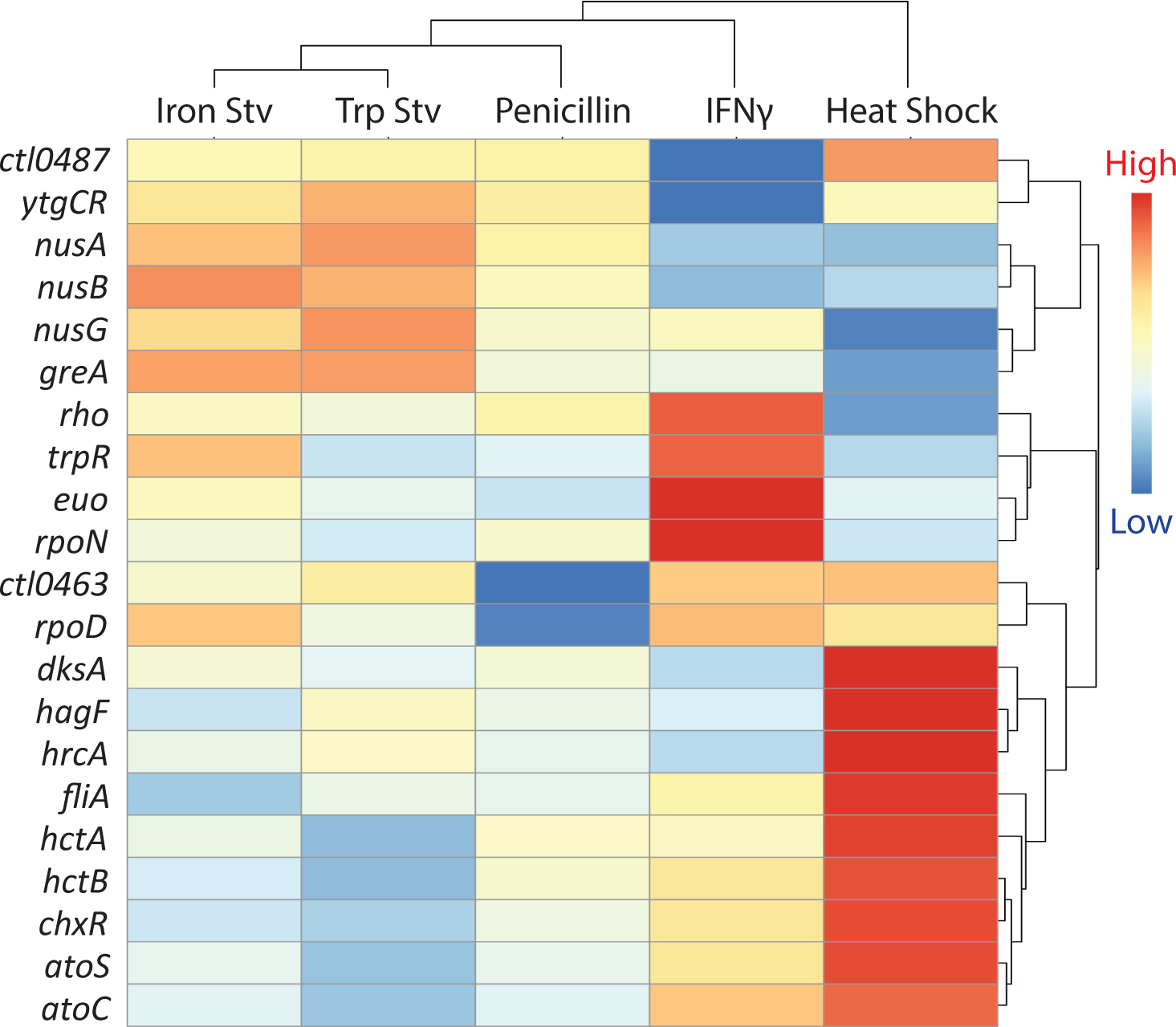
Stress-dependent expression patterns of transcriptional regulators. Hierarchical clustering heatmap of transcriptional regulator genes that are differentially expressed in at least one stress transcriptome. Clustering, dendrogram interpretation, and color scaling are as described in Fig. 1. Each row represents a transcriptional regulator, and each column represents a stress condition.

A distinct heat shock-associated cluster comprised regulators that were strongly and selectively induced in response to heat shock. This group included *hrcA*, the heat-inducible repressor of chaperone genes (51–53); *hagF*, a recently identified heat-responsive antagonist of *hrcA* (53); and the late developmental histone-like genes *hctA* and *hctB* (54, 55). In addition, multiple elongation factors (*nusA, nusB,* and *nusG*) were upregulated. Together, these regulators define a coordinated heat shock-specific transcriptional state characterized by enhanced transcriptional processivity and protein quality control.

In contrast, a chronic stress-associated cluster was defined by regulators that were consistently elevated across IFNγ, tryptophan starvation, iron starvation, and penicillin treatment but not during heat shock (Fig. 4). This group included *euo*, a repressor of late developmental gene expression (56, 57), as well as nutrient-responsive regulators such as *trpR* and *ytgR*. Their shared induction is consistent with sustained metabolic restriction and suppression of late developmental programs during stresses (58, 59).

Beyond these two dominant clusters, several regulators displayed stress-specific or heterogeneous expression patterns. Core sigma factor genes *rpoD* (σ66)*, rpoN* (σ54), and *fliA* (σ28) varied across chronic stress conditions rather than showing uniform regulation. Such differential regulation likely underlies features such as the partial divergence between IFNγ and tryptophan starvation transcriptomes, despite their shared impact on amino acid availability.

### Genes commonly regulated in all five stress transcriptomes

Despite the extensive stress-specific transcriptional divergence described above, we next asked whether any genes were consistently regulated across multiple stress transcriptomes. The numbers of genes uniquely or jointly altered in the five stress transcriptomes were compared using Venn diagrams (Fig. 5). The four chronic stress transcriptomes shared 20 upregulated and 99 downregulated genes. When the heat shock transcriptome was included in the comparison, the number of commonly upregulated and commonly downregulated genes decreased to four in each group (Fig. 5).

**Fig. 5.**
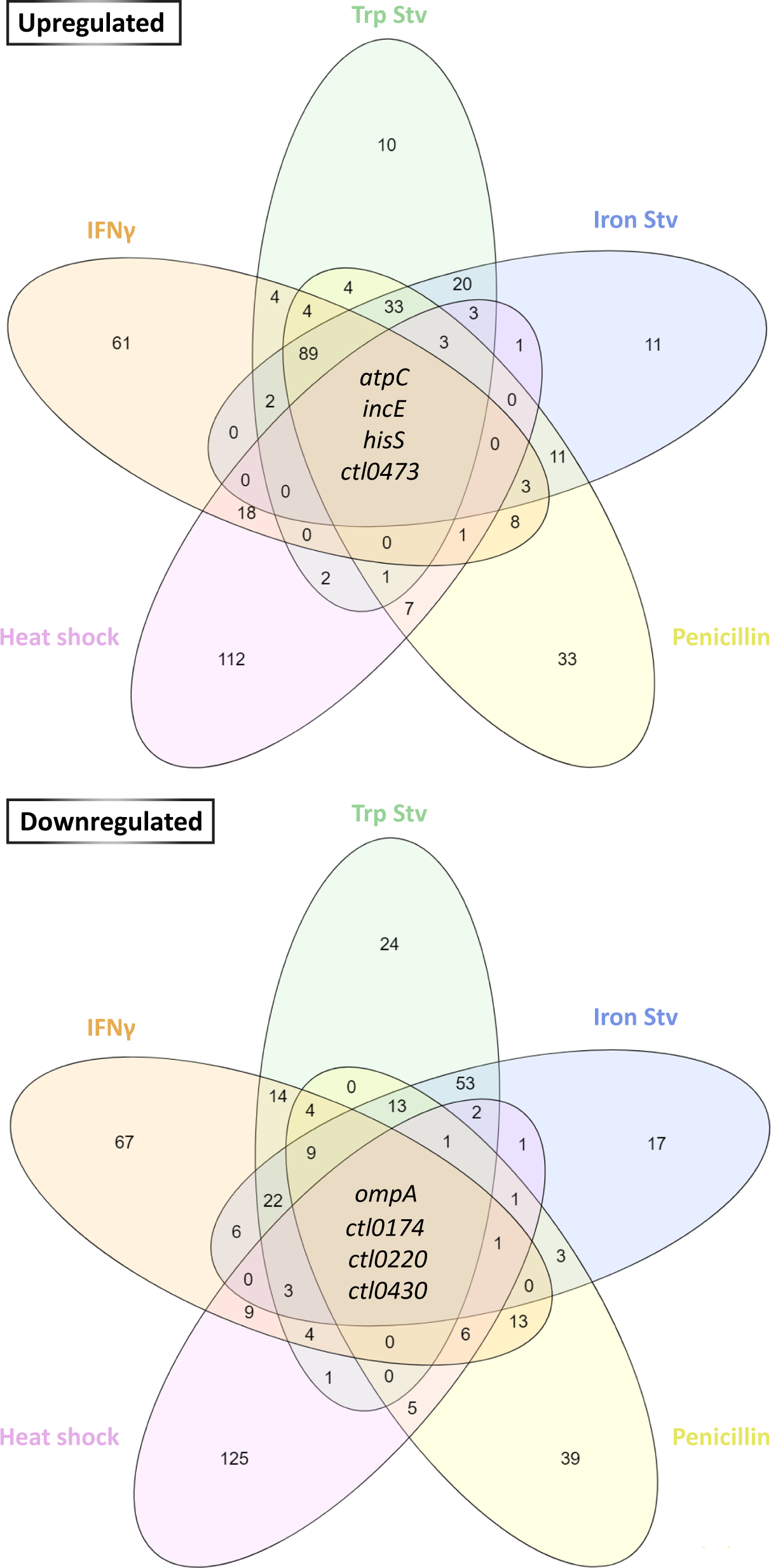
Genes commonly regulated across five stress transcriptomes. Venn diagrams show the numbers of overlapping upregulated (top) and downregulated genes (bottom) among stress transcriptomes, based on differential expression criteria applied to each transcriptome. The eight genes commonly upregulated across all five stress transcriptomes are identified.

Of the four genes commonly upregulated in all five stress transcriptomes, *atpC*, *hisS*, *incE*, and *ctl0473/ct221* encode an ATP synthase subunit, histidine-tRNA ligase, inclusion membrane protein E, and a hypothetical protein, respectively. Among the four commonly downregulated genes, *ompA* and *ctl0220/ct848* encode the major outer membrane protein (MOMP) and a T3SS effector, respectively; the remaining two genes, *ctl0174/ct805* and *ctl0430/ct178,* encode hypothetical proteins. The consistent regulation of these genes across all stress conditions suggests their involvement in stress responses, either as active drivers of adaptation or as passive markers of shared physiological states.

## Discussion

Stress responses are central to chlamydial pathophysiology (11–20, 60, 61). By comparing transcriptomes derived from cultures exposed to IFNγ, tryptophan starvation, iron starvation, penicillin, or heat shock, this study provides an integrated view of how *C. trachomatis* reprograms gene expression in response to diverse adverse conditions. This comparative analysis reveals that *C. trachomatis* employs distinct transcriptional programs that reflect the nature of the biological challenge it encounters.

At a broad level, the transcriptomic relatedness among stress conditions reflects fundamental differences in the nature of the stress imposed. Consistent with this distinction, hierarchical clustering and correlation analyses reveal a clear separation between the heat shock transcriptome and the remaining stress transcriptomes. The experimental procedures used to model the heat shock response represent an acute perturbation that rapidly disrupts protein homeostasis, whereas IFNγ treatment, tryptophan starvation, iron starvation, and penicillin exposure impose prolonged limitations on growth-associated biosynthetic capacity.

One notable feature of the chronic stress transcriptomes is the broad upregulation of ribosomal and protein synthesis genes, a pattern that contrasts with canonical bacterial stress responses (62). This behavior may be related to the absence of the *rel* gene in *Chlamydia*. In free-living bacteria, similar transcriptional profiles are typically observed only in starved *rel*-deficient mutants (2, 63, 64). Rel synthesizes the alarmone (p)ppGpp in response to amino acid starvation and other stressors (2, 65). The binding of (p)ppGpp to RNA polymerase leads to the repression of ribosomal and translational genes (2, 63, 64). Because *rel*-deficient mutants are generally more vulnerable under stress conditions (2, 63, 64), how elevated expression of ribosomal and protein synthesis genes contributes to chlamydial fitness during persistence remains an important unresolved question.

Within the group of chronic stresses, however, transcriptomic relatedness does not follow expectations based on the nominal classification of stressors, particularly the expectation that IFNγ treatment would most closely resemble tryptophan starvation. Instead, the close resemblance between tryptophan starvation and iron starvation is more pronounced than their resemblance to IFNγ treatment. This unexpected relationship can be explained by regulatory cross-talk centered on the iron-dependent transcriptional repressor YtgR (58, 59, 66). In *C. trachomatis*, translation of YtgR is gated by a tryptophan-rich regulatory motif, such that tryptophan limitation suppresses YtgR synthesis, whereas iron starvation compromises YtgR repressor activity by limiting availability of the iron cofactor required for DNA binding and repression (58, 59). As a result, both stresses converge on reduced YtgR function and a shared downstream regulatory state, providing a mechanistic basis for their close similarity.

By comparison, IFNγ treatment shows only partial overlap with tryptophan starvation. Although IFNγ-induced tryptophan depletion via host indoleamine 2,3-dioxygenase is a well-established antichlamydial mechanism, IFNγ signaling also activates additional antimicrobial programs in human cells that are not recapitulated by tryptophan deprivation alone, including alterations in host iron handling, metabolic reprogramming, and other cell-autonomous defenses. Supporting this, human genetic and cellular studies demonstrate that IFNγ retains substantial antimicrobial activity even when tryptophan depletion is attenuated (67). Therefore, the intermediate positioning of the IFNγ transcriptome likely reflects its multifactorial nature as a host-imposed stress that combines biosynthetic limitation with additional pressures beyond tryptophan starvation (22, 25).

Despite extensive stress-specific transcriptional reprogramming, overlap exists among transcriptomes induced by chronic stress conditions. The four chronic stress transcriptomes share approximately 120 commonly regulated genes, whereas inclusion of the heat shock transcriptome reduces this overlap to only eight genes. This graded pattern indicates that transcriptional convergence in *C. trachomatis* is stressor-dependent. Importantly, the biological significance of commonly regulated genes may differ. Consistently upregulated genes are more likely to serve functions actively involved in adaptation or survival under persistence-inducing environments, whereas commonly downregulated genes more plausibly reflect secondary consequences of growth and developmental defects.

In this context, the identification of stress-responsive pathways has implications for therapeutic intervention. Several genes that are consistently upregulated across chronic stress conditions encode functions that have been validated as antimicrobial targets in other bacterial systems. For example, bacterial tRNA synthetases are established drug targets, as illustrated by mupirocin, which inhibits isoleucyl-tRNA synthetase and is used to treat staphylococcal infections (68). Similarly, bacterial ATP synthase represents a proven vulnerability in persistent pathogens; bedaquiline targets the F₀F₁ ATP synthase of *Mycobacterium tuberculosis* and is highly effective against drug-resistant and nonreplicating bacilli (69, 70). Together, these precedents support the idea that targeting functions that are consistently upregulated and involved across multiple stress-induced transcriptional states may represent a viable strategy for disrupting chlamydial persistence.

In conclusion, our findings demonstrate that adaptation to different biological stressors in *C. trachomatis* is driven by distinct transcriptomic reprogramming, while consistently involving a set of commonly regulated genes. The products of these genes may represent shared points of vulnerability across persistence-inducing environments. This work has implications for the development of future antichlamydial strategies targeting chlamydial persistence.

## MATERIALS AND METHODS

### RNA-Seq datasets and experimental conditions

Stress conditions and corresponding NCBI accession numbers for the raw RNA-Seq datasets are listed in Table 1.

### Analysis of RNA-Seq data

Raw RNA-Seq data were downloaded from the NCBI repositories listed in Table 1 and reprocessed using a unified workflow on Galaxy (71). Adapter sequences were trimmed, and low-quality reads were removed using Trimmomatic (version 0.38). Reads were aligned to either the *C. trachomatis* serovar D genome (strain UW-3/CX; chromosome: GCF_000008725.1, plasmid: NC_020986.1) or the serovar L2 genome (strain 434/Bu; chromosome: GCF_000068585.1_ASM6858v1, plasmid: AM886278) using HISAT2 (version 2.2.1). Gene expression was quantified using featureCounts (version 2.0.3) to obtain raw read counts per gene. Differential expression was assessed using DESeq2 (version 2.11.40.8) with Benjamini–Hochberg correction for multiple testing (72, 73). Genes with an adjusted *P* value (Padj) < 0.05 and a fold change ≥ 1.5 (log2FC ≥ 0.58496) were considered differentially expressed.

### Annotation of hypothetical protein genes and their ontology assignments

Genes annotated as “hypothetical protein” in the reference genomes were evaluated to refine functional descriptions and to support ontology assignments used in the functional-category analyses. For each hypothetical protein gene, we queried ChlamBase to retrieve curated locus information, alternative gene names, and any community- or literature-derived annotations for the corresponding gene product (74). We also reviewed the corresponding UniProtKB entry to extract the current protein name, description, predicted features, and evidence context for functional annotations (75).

To incorporate experimentally supported information not captured in database summaries, we performed targeted PubMed searches using locus tags and commonly used aliases (e.g., CT_694, CTL_0360), as well as gene and protein name synonyms. When peer-reviewed studies provided direct experimental evidence (e.g., secretion via the type III secretion system, inclusion membrane localization, interaction partners, or phenotypes associated with targeted mutagenesis or knockdown), we used those findings to refine the functional description applied in this study and to guide assignment to the appropriate ontology category (e.g., type III secretion effectors, inclusion membrane proteins, or general function prediction).

## Figure preparation

Hierarchical clustering heatmaps were generated using the R package pheatmap (76). Pearson correlation analysis was conducted in Microsoft Excel. Divergent bar graphs were created in GraphPad Prism. Venn diagrams were generated using the web-based visualization tool InteractiVenn (77).

## ACKNOWLEDGMENTS

This work was supported by the National Institutes of Health grant numbers AI140167 and AI154305 (to H.F.) and AI182210 (to G.Z. and H.F.).

## REFERENCES

1. Harms A, Maisonneuve E, Gerdes K. 2016. Mechanisms of bacterial persistence during stress and antibiotic exposure. Science 354.

2. Das B, Bhadra RK. 2020. (p)ppGpp Metabolism and Antimicrobial Resistance in Bacterial Pathogens. Front Microbiol 11:563944.

3. Gollan B, Grabe G, Michaux C, Helaine S. 2019. Bacterial Persisters and Infection: Past, Present, and Progressing. Annu Rev Microbiol 73:359–385.

4. Elwell C, Mirrashidi K, Engel J. 2016. Chlamydia cell biology and pathogenesis. Nat Rev Microbiol 14:385–400.

5. Zhong G, Brunham RC, de la Maza LM, Darville T, Deal C. 2017. National Institute of Allergy and Infectious Diseases workshop report: “*Chlamydia* vaccines: the way forward”. Vaccine doi:doi.org/10.1016/j.vaccine.2017.10.075.

6. McCullough A, Steven H, and Weber MM. 2025. Pathogenicity and virulence of Chlamydia trachomatis: Insights into host interactions, immune evasion, and intracellular survival. Virulence 16:2503423.

7. López-Hurtado M, Escarcega-Tame MA, Escobedo-Guerra MR, de Haro-Cruz MJ, Guerra-Infante FM. 2022. Identification of Chlamydia trachomatis genotypes in Mexican men with infertile women as sexual partners. Enferm Infecc Microbiol Clin (Engl Ed) 40:353–358.

8. Bryan ER, McLachlan RI, Rombauts L, Katz DJ, Yazdani A, Bogoevski K, Chang C, Giles ML, Carey AJ, Armitage CW, Trim LK, McLaughlin EA, Beagley KW. 2019. Detection of chlamydia infection within human testicular biopsies. Human Reproduction 34:1891–1898.

9. Hybiske K, Stephens RS. 2007. Mechanisms of *Chlamydia trachomatis* entry into nonphagocytic cells. Infect Immun 75:3925–3934.

10. Hybiske K, Stephens RS. 2007. Mechanisms of host cell exit by the intracellular bacterium *Chlamydia*. Proc Natl Acad Sci USA 104:11430–11435.

11. Belland RJ, Nelson DE, Virok D, Crane DD, Hogan D, Sturdevant D, Beatty WL, Caldwell HD. 2003. Transcriptome analysis of chlamydial growth during IFN-γ-mediated persistence and reactivation. Proc Natl Acad Sci USA 100:15971–15976.

12. Yang C, Kari L, Lei L, Carlson JH, Ma L, Couch CE, Whitmire WM, Bock K, Moore I, Bonner C, McClarty G, Caldwell HD. 2020. Chlamydia trachomatis Plasmid Gene Protein 3 Is Essential for the Establishment of Persistent Infection and Associated Immunopathology. mBio 11.

13. Brinkworth AJ, Wildung MR, Carabeo RA. 2018. Genomewide Transcriptional Responses of Iron-Starved Chlamydia trachomatis Reveal Prioritization of Metabolic Precursor Synthesis over Protein Translation. mSystems 3:e00184–17.

14. Huston WM, Theodoropoulos C, Mathews SA, Timms P. 2008. Chlamydia trachomatis responds to heat shock, penicillin induced persistence, and IFN-gamma persistence by altering levels of the extracytoplasmic stress response protease HtrA. BMC Microbiol 8:190.

15. Brockett MR, Liechti GW. 2021. Persistence Alters the Interaction between Chlamydia trachomatis and Its Host Cell. Infect Immun 89:e0068520.

16. Shima K, Kaufhold I, Eder T, Kading N, Schmidt N, Ogunsulire IM, Deenen R, Kohrer K, Friedrich D, Isay SE, Grebien F, Klinger M, Richer BC, Gunther UL, Deepe GS, Jr., Rattei T, Rupp J. 2021. Regulation of the mitochondrion-fatty acid axis for the metabolic reprogramming of *Chlamydia trachomatis* during treatment with beta-lactam antimicrobials. mBio 12.

17. Panzetta ME, Valdivia RH, Saka HA. 2018. Chlamydia Persistence: A Survival Strategy to Evade Antimicrobial Effects in-vitro and in-vivo. Frontiers in Microbiology 9.

18. Rockey DD, Wang X, Debrine A, Grieshaber N, Grieshaber SS. 2024. Metabolic dormancy in Chlamydia trachomatis treated with different antibiotics. Infect Immun 92:e0033923.

19. Kozusnik T, Adams SE, Greub G. 2024. Aberrant Bodies: An Alternative Metabolic Homeostasis Allowing Survivability? Microorganisms 12.

20. Jury B, Fleming C, Huston WM, Luu LDW. 2023. Molecular pathogenesis of Chlamydia trachomatis. Front Cell Infect Microbiol 13:1281823.

21. Huang Y, Wurihan W, Lu B, Zou Y, Wang Y, Weldon K, Fondell JD, Lai Z, Wu X, Fan H. 2021. Robust heat shock response in *Chlamydia* lacking a typical heat shock sigma factor. Front Microbiol 12:812448.

22. Muramatsu MK, Brothwell JA, Stein BD, Putman TE, Rockey DD, Nelson DE. 2016. Beyond Tryptophan Synthase: Identification of Genes That Contribute to Chlamydia trachomatis Survival during Gamma Interferon-Induced Persistence and Reactivation. Infect Immun 84:2791–801.

23. Olaleye AO, Babah OA, Osuagwu CS, Ogunsola FT, Afolabi BB. 2020. Sexually transmitted infections in pregnancy – An update on *Chlamydia trachomatis* and *Neisseria gonorrhoeae*. European Journal of Obstetrics and Gynecology and Reproductive Biology 255:1–12.

24. Pokorzynski ND, Alla MR, Carabeo RA. 2022. Host Cell Amplification of Nutritional Stress Contributes To Persistence in Chlamydia trachomatis. mBio doi:10.1128/mbio.02719-22:e0271922.

25. Reitano JR, Coers J. 2024. Restriction and evasion: a review of IFNγ-mediated cell-autonomous defense pathways during genital Chlamydia infection. Pathog Dis 82.

26. Nsereko E, Uwase A, Mukabutera A, Muvunyi CM, Rulisa S, Ntirushwa D, Moreland P, Corwin EJ, Santos N, Nzayirambaho M, Wojcicki JM. 2020. Maternal genitourinary infections and poor nutritional status increase risk of preterm birth in Gasabo District, Rwanda: a prospective, longitudinal, cohort study. BMC Pregnancy Childbirth 20:345.

27. Ray A, Moore TF, Naik DSL, Borsch DM. 2024. Insights into the Two Most Common Cancers of Primitive Gut-Derived Structures and Their Microbial Connections. Medicina (Kaunas) 60.

28. Chudomirova K, Abadjieva T, Yankova R. 2008. Clinical tetrad of arthritis, urethritis, conjunctivitis, and mucocutaneous lesions (HLA-B27-associated spondyloarthropathy, Reiter syndrome): report of a case. Dermatol Online J 14:4.

29. Leslie SW, Vinod J. 2025. Lymphogranuloma Venereum Infection, StatPearls. StatPearls Publishing Copyright © 2025, StatPearls Publishing LLC., Treasure Island (FL).

30. Wu L, Chen L, Peng L, Liu C, He S, Xie L. 2025. Clinical characteristics of Chlamydia psittaci pneumonia and predictors analysis of severe patients: a retrospective observational study. Front Med (Lausanne) 12:1565254.

31. Rohde G, Straube E, Essig A, Reinhold P, Sachse K. 2010. Chlamydial Zoonoses. Deutsches Arzteblatt International 107:174–180.

32. de Voux A, Kent JB, Macomber K, Krzanowski K, Jackson D, Starr T, Johnson S, Richmond D, Crane LR, Cohn J, Finch C, McFadden J, Pillay A, Chen C, Anderson L, Kersh EN. 2016. Notes from the Field: Cluster of Lymphogranuloma Venereum Cases Among Men Who Have Sex with Men - Michigan, August 2015-April 2016. MMWR Morb Mortal Wkly Rep 65:920–1.

33. Dan M, Tyrrell LD, Goldsand G. 1987. Isolation of Chlamydia trachomatis from the liver of a patient with prolonged fever. Gut 28:1514–6.

34. Narberhaus F. 1999. Negative regulation of bacterial heat shock genes. Molecular Microbiology 31:1–8.

35. Gragerov A, Nudler E, Komissarova N, Gaitanaris GA, Gottesman ME, Nikiforov V. 1992. Cooperation of GroEL/GroES and DnaK/DnaJ heat shock proteins in preventing protein misfolding in Escherichia coli. Proc Natl Acad Sci U S A 89:10341–4.

36. Lu B, Wang Y, Wurihan W, Cheng A, Yeung S, Fondell JD, Lai Z, Wan D, Wu X, Li WV, Fan H. 2024. Requirement of GrgA for *Chlamydia* infectious progeny production, optimal growth, and efficient plasmid maintenance. mBio 15:e0203623.

37. Hatch ND, Ouellette SP. 2023. Identification of the alternative sigma factor regulons of *Chlamydia trachomatis* using multiplexed CRISPR interference. mSphere doi:10.1128/msphere.00391-23:e0039123.

38. Soules KR, LaBrie SD, May BH, Hefty PS. 2020. Sigma 54-regulated transcription is associated with membrane reorganization and type III secretion effectors during conversion to infectious forms of *Chlamydia trachomatis*. mBio 11.

39. Rucks EA. 2023. Type III Secretion in Chlamydia. Microbiol Mol Biol Rev 87:e0003423.

40. Lutter EI, Barger AC, Nair V, Hackstadt T. 2013. Chlamydia trachomatis inclusion membrane protein CT228 recruits elements of the myosin phosphatase pathway to regulate release mechanisms. Cell reports 3:1921–31.

41. Yang C, Starr T, Song L, Carlson JH, Sturdevant GL, Beare PA, Whitmire WM, Caldwell HD. 2015. Chlamydial Lytic Exit from Host Cells Is Plasmid Regulated. mBio 6:e01648–15.

42. Nguyen PH, Lutter EI, Hackstadt T. 2018. Chlamydia trachomatis inclusion membrane protein MrcA interacts with the inositol 1,4,5-trisphosphate receptor type 3 (ITPR3) to regulate extrusion formation. PLoS Pathog 14:e1006911.

43. Pereira IS, Pais SV, Borges V, Borrego MJ, Gomes JP, Mota LJ. 2022. The Type III Secretion Effector CteG Mediates Host Cell Lytic Exit of Chlamydia trachomatis. Front Cell Infect Microbiol 12:902210.

44. Liu X, Afrane M, Clemmer DE, Zhong G, Nelson DE. 2010. Identification of Chlamydia trachomatis outer membrane complex proteins by differential proteomics. J Bacteriol 192:2852–60.

45. Zhang T, Huo Z, Ma J, He C, Zhong G. 2019. The Plasmid-Encoded pGP3 Promotes Chlamydia Evasion of Acidic Barriers in Both Stomach and Vagina. Infect Immun 87.

46. Liu Y, Huang Y, Yang Z, Sun Y, Gong S, Hou S, Chen C, Li Z, Liu Q, Wu Y, Baseman J, Zhong G. 2014. Plasmid-encoded Pgp3 is a major virulence factor for Chlamydia muridarum to induce hydrosalpinx in mice. Infect Immun 82:5327–35.

47. Zhong G. 2017. Chlamydial plasmid-dependent pathogenicity. Trends in microbiology 25:141–152.

48. Turman BJ, Darville T, O’Connell CM. 2023. Plasmid-mediated virulence in Chlamydia. Front Cell Infect Microbiol 13:1251135.

49. Song L, Carlson JH, Whitmire WM, Kari L, Virtaneva K, Sturdevant DE, Watkins H, Zhou B, Sturdevant GL, Porcella SF, McClarty G, Caldwell HD. 2013. *Chlamydia trachomatis* plasmid-encoded Pgp4 is a transcriptional regulator of virulence-associated genes. Infect Immun 81:636–644.

50. Zhang Q, Rosario CJ, Sheehan LM, Rizvi SM, Brothwell JA, He C, Tan M. 2020. The repressor function of the *Chlamydia* late regulator EUO is enhanced by the plasmid-encoded protein Pgp4. Journal of Bacteriology 202:e00793–19.

51. Wilson AC, Wu CC, Yates JR, 3rd, Tan M. 2005. Chlamydial GroEL autoregulates its own expression through direct interactions with the HrcA repressor protein. J Bacteriol 187:7535–42.

52. Wilson AC, Tan M. 2004. Stress response gene regulation in Chlamydia is dependent on HrcA-CIRCE interactions. J Bacteriol 186:3384–91.

53. Huang Y, Wang Y, Pan M, Wan D, Wang L, Fondell JD, Wu X, Zhong G, Fan H. 2025. A lineage-specific heat-induced feedback loop controls HrcA to promote chlamydial fitness under stress. bioRxiv doi:10.1101/2025.05.30.657042.

54. Belland RJ, Zhong G, Crane DD, Hogan D, Sturdevant D, Sharma J, Beatty WL, Caldwell HD. 2003. Genomic transcriptional profiling of the developmental cycle of *Chlamydia trachomatis*. Proc Natl Acad Sci USA 100:8478–83.

55. Chiarelli TJ, Grieshaber NA, Omsland A, Remien CH, Grieshaber SS. 2020. Single-Inclusion Kinetics of Chlamydia trachomatis Development. mSystems 5.

56. Hakiem OR, Rizvi SMA, Ramirez C, Tan M. 2023. Euo is a developmental regulator that represses late genes and activates midcycle genes in Chlamydia trachomatis. mBio 14:e0046523.

57. Rosario CJ, Hanson BR, Tan M. 2014. The transcriptional repressor EUO regulates both subsets of Chlamydia late genes. Mol Microbiol 94:888–97.

58. Pokorzynski ND, Brinkworth AJ, Carabeo R. 2019. A bipartite iron-dependent transcriptional regulation of the tryptophan salvage pathway in Chlamydia trachomatis. eLife 8:e42295.

59. Pokorzynski ND, Hatch ND, Ouellette SP, Carabeo RA. 2020. The iron-dependent repressor YtgR is a tryptophan-dependent attenuator of the trpRBA operon in Chlamydia trachomatis. Nat Commun 11:6430.

60. Mirrashidi KM, Elwell CA, Verschueren E, Johnson JR, Frando A, Von Dollen J, Rosenberg O, Gulbahce N, Jang G, Johnson T, Jager S, Gopalakrishnan AM, Sherry J, Dunn JD, Olive A, Penn B, Shales M, Cox JS, Starnbach MN, Derre I, Valdivia R, Krogan NJ, Engel J. 2015. Global mapping of the Inc-human interactome reveals that retromer restricts Chlamydia infection. Cell Host Microbe 18:109–21.

61. Olson-Wood MG, Jorgenson LM, Ouellette SP, Rucks EA. 2021. Inclusion membrane growth and composition are altered by overexpression of specific inclusion membrane proteins in Chlamydia trachomatis L2. Infect Immun 89:e0009421.

62. Durfee T, Hansen AM, Zhi H, Blattner FR, Jin DJ. 2008. Transcription profiling of the stringent response in Escherichia coli. J Bacteriol 190:1084–96.

63. Goodman D. 1970. Ribosomal protein synthesis during amino acid starvation and chloramphenicol treatment. Journal of Molecular Biology 51:491–499.

64. Dalebroux ZD, Swanson MS. 2012. ppGpp: magic beyond RNA polymerase. Nat Rev Microbiol 10:203–12.

65. Gratani FL, Horvatek P, Geiger T, Borisova M, Mayer C, Grin I, Wagner S, Steinchen W, Bange G, Velic A, Maček B, Wolz C. 2018. Regulation of the opposing (p)ppGpp synthetase and hydrolase activities in a bifunctional RelA/SpoT homologue from Staphylococcus aureus. PLOS Genetics 14:e1007514.

66. Akers JC, HoDac H, Lathrop RH, Tan M. 2011. Identification and functional analysis of CT069 as a novel transcriptional regulator in *Chlamydia*. Journal of Bacteriology 193:6123–6131.

67. Murray HW, Szuro-Sudol A, Wellner D, Oca MJ, Granger AM, Libby DM, Rothermel CD, Rubin BY. 1989. Role of tryptophan degradation in respiratory burst-independent antimicrobial activity of gamma interferon-stimulated human macrophages. Infect Immun 57:845–9.

68. Simmons KJ, Chopra I, Fishwick CW. 2010. Structure-based discovery of antibacterial drugs. Nat Rev Microbiol 8:501–10.

69. Melander RJ, Zurawski DV, Melander C. 2018. Narrow-Spectrum Antibacterial Agents. Medchemcomm 9:12–21.

70. Vestergaard M, Bald D, Ingmer H. 2022. Targeting the ATP synthase in bacterial and fungal pathogens: beyond Mycobacterium tuberculosis. J Glob Antimicrob Resist 29:29–41.

71. Community TG. 2024. The Galaxy platform for accessible, reproducible, and collaborative data analyses: 2024 update. Nucleic Acids Research 52:W83–W94.

72. Love M, Anders S, Huber W. 2014. Differential analysis of count data the DESeq2 package. Genome Biol 15:1–41.

73. Love MI, Huber W, Anders S. 2014. Moderated estimation of fold change and dispersion for RNA-seq data with DESeq2. Genome Biol 15:550.

74. Putman T, Hybiske K, Jow D, Afrasiabi C, Lelong S, Cano MA, Stupp GS, Waagmeester A, Good BM, Wu C, Su AI. 2019. ChlamBase: a curated model organism database for the Chlamydia research community. Database (Oxford) 2019.

75. Consortium TU. 2020. UniProt: the universal protein knowledgebase in 2021. Nucleic Acids Research 49:D480–D489.

76. Kolde R. 2010. pheatmap: Pretty Heatmaps. doi:10.32614/CRAN.package.pheatmap.

77. Heberle H, Meirelles GV, da Silva FR, Telles GP, Minghim R. 2015. InteractiVenn: a web-based tool for the analysis of sets through Venn diagrams. BMC Bioinformatics 16:169.

