## Supplementary figures and images for "Comparative Transcriptomics Reveals Genes Commonly Induced by Distinct Stressors in *Chlamydia*"

### Fig. S1

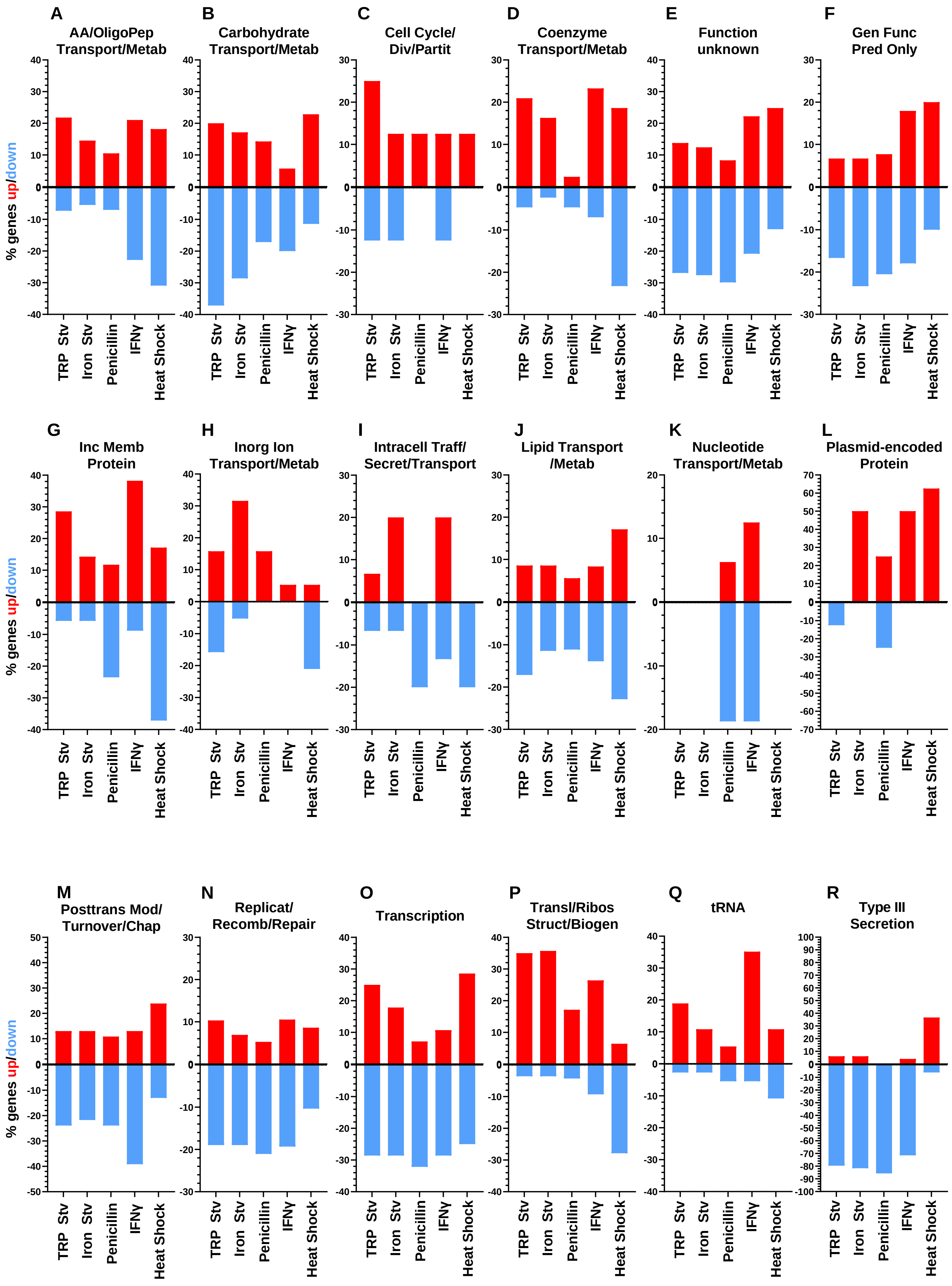
